# Spatial transcriptomics unveils landscape of resistance to concurrent chemo-radiotherapy in hypopharyngeal squamous cell carcinoma: the role of *SPP1*^+^ macrophages

**DOI:** 10.1101/2024.07.09.602476

**Authors:** Jungyoon Ohn, Sungwoo Bae, Hongyoon Choi, In Gul Kim, Kwon Joong Na, Eun-Jae Chung

**Author notes:** **Correspondence and Reprint request Kwon Joong Na, MD**, Portrai, Inc, Dongsunragil, 78-18, Jongno-gu, Seoul, Republic of Korea (03136), Department of Thoracic and Cardiovascular Surgery, Seoul National University Hospital, 101, Daehak-ro, Jongno-gu, Seoul, Republic of Korea (03080), **Eun-Jae Chung, MD, PhD**, Department of Otolaryngology-Head and Neck Surgery, Seoul National University College of Medicine, 101 Daehak-ro, Jongno-gu, Seoul, Republic of Korea (03080). These authors contributed equally to this work (Jungyoon Ohn, Sungwoo Bae).

## Abstract

Hypopharyngeal squamous cell carcinoma (SCC) is a highly aggressive cancer with a poor prognosis, particularly in advanced stages where concurrent chemoradiotherapy (CCRT) is used for treatment. However, resistance to CCRT poses a significant challenge, often leading to treatment failure and disease progression. This study explores the tumor microenvironment (TME) of hypopharyngeal SCC to understand the molecular mechanisms underlying CCRT resistance. Using spatial transcriptomics (ST), we analyzed tissue samples from patients with locally advanced hypopharyngeal SCC, distinguishing between those who were CCRT-resistant and those who were CCRT-naive. The analysis revealed six distinct cellular clusters within the TME, including a prominent epithelio-immune cellular area in CCRT-resistant tissues. SPP1 was identified as a key gene with significantly higher expression in CCRT-resistant samples, specifically within macrophages. Further investigation showed that SPP1+ macrophages interacted with malignant epithelial cells through SPP1-CD44 and SPP1-ITGB1 ligand-receptor pairs. These interactions were primarily localized in the peri-tumoral and intra-tumoral regions, highlighting their potential role in driving CCRT resistance. Our findings suggest that SPP1+ macrophages contribute to the resistant phenotype in hypopharyngeal SCC by modulating the TME and interacting with cancer cells. Understanding these interactions offers valuable insights into the mechanisms of CCRT resistance and may inform the development of targeted therapies to improve patient outcomes.

## Main text

Hypopharyngeal squamous cell carcinoma (SCC) is a highly aggressive cancer that is notorious for poor prognosis. Regardless of the poor prognosis, the most critical issue for patients and caregivers in the treatment of hypopharyngeal SCC is whether the larynx can be preserved to maintain normal breathing, speaking, and eating functions. Laryngeal preservation through surgical treatment is only possible in early-stage cancer. Advanced tumors of the hypopharynx (over T3) are mostly managed by total laryngectomy followed by radiation therapy or concurrent chemoradiotherapy (CCRT) called organ preservation protocol. The organ preservation protocol can preserve the larynx. However, there is currently no method to predict the CCRT resistance on the individual patient. [1, 2]

Despite the established efficacy of CCRT, the clinical landscape is frequently marred by the emergence of CCRT resistance, precipitating therapeutic failure and progression of disease. Critically, CCRT exerts multifaceted influences on the immune and stromal cellular constituents of the tumor microenvironment (TME). [3] This modulation of the TME can subsequently alter the biological aggressiveness of the tumor, underscoring the complexity of treatment response and resistance in hypopharyngeal SCC. Consequently, to surmount CCRT resistance and identify novel biomarkers predictive of such resistance, a profound understanding of the TME dynamics under these therapeutic conditions is paramount. [3] This study aims to explore the tumor microenvironment (TME) of hypopharyngeal SCC which recur after CCRT, with the objective of identifying spatial molecular signatures within the cancer tissue using spatial transcriptomics (ST). By employing ST, we seek to gain insights into the spatial distribution of transcriptomic changes associated with CCRT resistant in hypopharyngeal SCC. Such insights might hold promise for the development of targeted therapeutic strategies and improved patient management in CCRT resistant hypopharyngeal SCC.

Firstly, we obtained tissue samples from two patients with locally advanced hypopharyngeal SCC who had undergone CCRT and subsequently recurred (designated as CCRT-resistant) and from two patients with locally advanced hypopharyngeal SCC who had upfront surgery without CCRT (designated as CCRT-naive) (**Fig. 1A and Supplementary Table**). The tissues were processed to acquire H&E staining tissue image, and ST data of the tissues were obtained using the Visium platform (10x Genomics, USA). The acquired ST data were analyzed using bioinformatics tools (**Supplementary Method**). In both the CCRT-resistant and naive groups, H&E images showed the presence of hypopharyngeal SCC (**Fig. 1B**). Upon plotting the obtained ST data spots on the uniform manifold approximation and projection for dimension reduction (UMAP) plot, the spots from the four different tissues showed significant overlap, implying minimized batch effects (**Fig. 1C**). In addition, these spots clustered into six distinct groups based on key gene expression identifiers (**Fig. 1D, Supplementary Fig. 1, and Supplementary File**): Hypopharyngeal SCC, Cancer-associated fibroblasts, Epithelial cells, Immune cells, Muscle cells, and Epithelio-immune cell areas.

**Figure 1.**
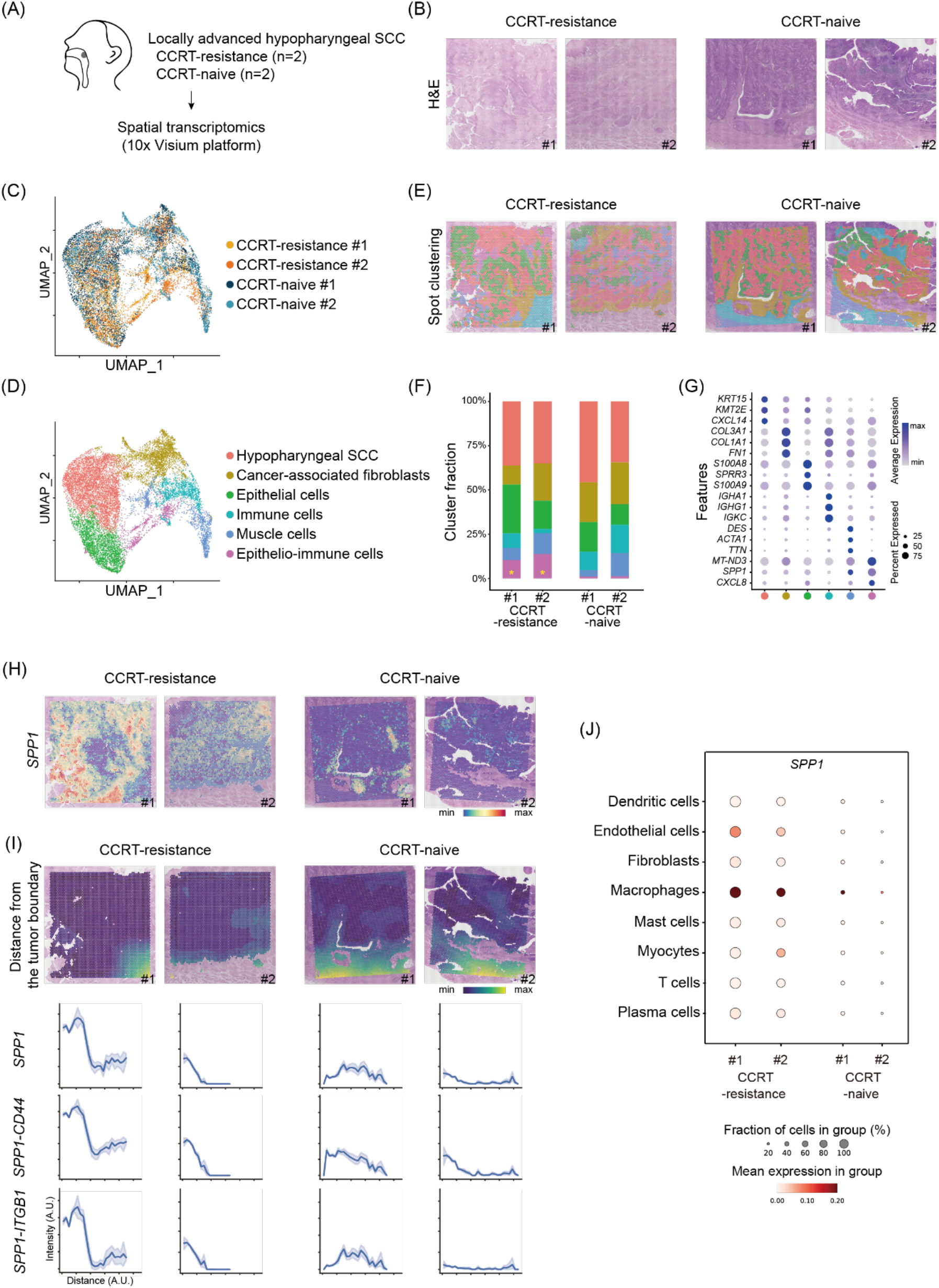
Intra- and peri-tumoral *SPP1+* macrophages are associated with CCRT-resistant hypopharyngeal SCC. **(A)** Schematic of spatial profiling for patients with hypopharyngeal SCC (n=2, CCRT-resistant; n=2, CCRT-naive). **(B)** H&E-stained tissue slides. **(C)** UMAP of ST spots color-coded by sample origin. **(D)** UMAP of ST spots color-coded by cluster identities defined based on their signature gene sets. **(E)** Spatial location of the spot clusters. **(F)** The fraction of the number of ST spots corresponding to the spot cluster in each sample. The yellow asterisk indicates a significant cluster in the CCRT-resistant group. **(G)** The top three signature genes of each cluster (selected with a log fold change threshold and minimum expression percentage both set to 0.25). **(H)** The distribution pattern of *SPP1* gene expression for each sample overlaid on the H&E slide. **(I)** Relative spatial distance from the tumor boundary and corresponding *SPP1* expression, *SPP1-CD44* and *SPP1-ITGB1* interaction strength. **(J)** Estimated *SPP1* expression levels in each cell type within the TME. **Abbreviation**: CCRT, concurrent chemo-radiotherapy; SCC, squamous cell carcinoma; ST, spatial transcriptomics; TME, tumor microenvironment; UMAP: uniform manifold approximation and projection

Interestingly, the epithelio-immune cellular area was more prominent in the CCRT-resistant tissue samples compared to the CCRT-naïve tissue samples (**Fig. 1F**). *SPP1* was identified as a signature gene with the highest differential expression between the epithelio-immune cluster and other clusters (**Fig. 1G**). In line with these findings, when examining the expression pattern of the *SPP1* gene across the whole tissue, it was notably elevated in the CCRT-resistant TME compared to the CCRT-naïve group (**Fig. 1H**).

Along with the observation of high expression of *SPP1* in the CCRT-resistant tissue samples, ligand-receptor (LR) interaction analysis using *stLearn* was performed on each tissue sample, revealing the top 50 LR interactions respectively (**Supplementary Fig. 2**). Among the interactions, *SPP1* ligand-mediated interactions (*SPP1-CD44* and *SPP1-ITGB1*) were commonly found in the CCRT-resistant group; however they were not observed in the CCRT-naïve group. Next, we explored the spatial distribution of the *SPP1-CD44* and *SPP1-ITGB1* interactions within TME. To accurately characterize the histological architecture of each hypopharyngeal SCC into cancer or surrounding stroma in TME, *Cancer-Finder* tool was used [4]. Also histopathology of H&E stained slide images were reviewed by a board-certified pathologist (**Supplementary Fig. 3**). We found that these interactions were primarily detected in the peri-tumoral spatial region, as well as in the intra-tumoral spatial area (**Fig. 1I** and **Supplementary Fig. 4**).

To further investigate which cells within the diverse cell populations constituting the hypopharyngeal SCC TME predominantly express the *SPP1* gene, we utilized the bioinformatic tool *Cell2location*. It revealed significant expression of *SPP1* in macrophages among various cell types (**Fig. 1J** and **Supplementary Fig. 5**). Then, we aimed to identify the *SPP1*-mediated cell-cell interaction pairs involving SPP1+ macrophages in CCRT-resistant hypopharyngeal SCC tissues using the *stLearn* tool. The analysis showed a higher number of interactions from macrophages to malignant epithelial cells in the CCRT-resistant tissues compared to the CCRT-naïve tissues. The conspicuous absence of macrophage-to-malignant cell interactions in the CCRT-naive group was noteworthy (**Supplementary Fig. 6**).

Herein, it was found that CCRT-resistant hypopharyngeal SCC was characterized by *SPP1+* macrophages in TME which interacts with malignant epithelial cells in intra-tumoral and peri-tumoral regions via *SPP1-CD44* and *SPP1-ITGB1. SPP1* gene encodes Secreted phosphoprotein 1 protein, also known as osteopontin. *SPP1* is a multifunctional integrin-binding glycoprotein, which are over-expressed in a variety of tumors playing pivotal role in controlling the cell signaling pathway for tumor progression and metastasis. [5]

Notably, the *SPP1* gene has been identified as a significant biomarker in hypopharyngeal SCC, [6] with its upregulated mRNA expression in tumor tissues correlating with poor overall survival of hypopharyngeal SCC patients. [7] Additionally, its role in promoting lymph node metastasis (LNM) in hypopharyngeal SCC has been proposed. [8] Analysis of large public datasets [9] suggested the role of interactions, *SPP1-CD44* and *SPP1-ITGB1*, in hypopharyngeal SCC progression. CD44 has also been associated with worse tumor grade/staging, and prognosis in pharyngeal and laryngeal cancers. [10] Patients with hypopharyngeal SCC who exhibit concurrent expression of ITGB1 and NOTCH 1 protein experience significantly poorer survival rates, potentially through the regulation of cancer stemness. [11]

Of particular, *SPP1*+ tumor-associated macrophage infiltration is associated with LNM and poor prognosis in hypopharyngeal SCC patients. [12] Also, macrophages defined by *SPP1* and *CXCL9* expression is strongly associated with hypopharyngeal SCC prognosis. [13] The infiltration of *SPP1*+ macrophages gradually increases with tumor progression, and their interaction with cancer cells gradually increases to reprogram malignant cells, leading to tumor upstaging and shaping of the desmoplastic microenvironment. [14]

These precedent findings provide strong support for our analysis results. However, since the spatial transcriptomics captures RNA expression in the unit of spot, which consists of a mixture of cells, further validation at the single-cell and the protein levels is required in future studies. Through this study, we have identified characteristics of TME in patients experiencing recurrence after CCRT treatment, showing heightened interaction between *SPP1*-positive macrophages and cancer cells via *CD44* or *ITGB1*. Understanding the spatial distribution and functional implications of *SPP1*-expressing macrophage interacting with cancer cells within the TME could provide valuable insights into the mechanisms underlying CCRT resistance in hypopharyngeal SCC. This may ultimately help to improve clinical decision-making, guiding the development of more effective therapeutic strategies for hypopharyngeal SCC patients.

## DATA AVAILABILITY

The dataset generated for this study can be accessed upon a reasonable request.

## SUPPLEMENTARY DATA

Supplementary Table

Supplementary Figures 1-6

Supplementary Methods

Supplementary File

Supplementary References

## ACKNOWLEDGMENTS

We thank all members of Portrai, Inc. for the discussion and technical support. We also thank Ho Sub Park, a board-certified pathologist, who helped the pathologic analysis of the H&E-stained image.

## FUNDING

This research was supported by the National Research Foundation of Korea (2020M3A9B6037195, and RS-2024-00357094), and the SNUH

Research Fund (2620210050).

## CONFLICTS OF INTEREST

H.C. and K.J.N. are the co-founders and shareholders of Portrai, Inc.

## ETHICS DECLARATION

The study protocol was reviewed by the Institutional Review Board of Seoul National University Hospital, approved as a minimal-risk retrospective study (approval date: 27/12/2023, approval number: H-2203-011-1304), and individual consent was waived.

