## Supplementary data for "Spatial transcriptomics unveils landscape of resistance to concurrent chemo-radiotherapy in hypopharyngeal squamous cell carcinoma: the role of *SPP1*^+^ macrophages"

**Supplementary Table.** The clinical characteristics of the patients

| | Age | Sex | CCRT regimen | T stage | N stage | Stage# | *p16* IHC staining |
|---|---|---|---|---|---|---|---|
| CCRT-resistance #1 | 58 | Male | Weekly CDDP + RT | 2 | 2c | 4A | Negative |
| CCRT-resistance #2 | 60 | Male | Weekly CDDP + RT | 4a | 3b | 4B | Negative |
| CCRT-naïve #1 | 60 | Male | NA | 3 | 3b | 4A | Negative |
| CCRT-naïve #2 | 72 | Female | NA | 3 | 3b | 4B | Negative |

#Clinical stage was based on AJCC 7th edition.

**Abbreviation**: CCRT, concurrent chemo-radiotherapy; CDDP, cisplatin; IHC, immunohistochemistry; NA, not applicable; RT, radiotherapy

.

### Supplementary Figures

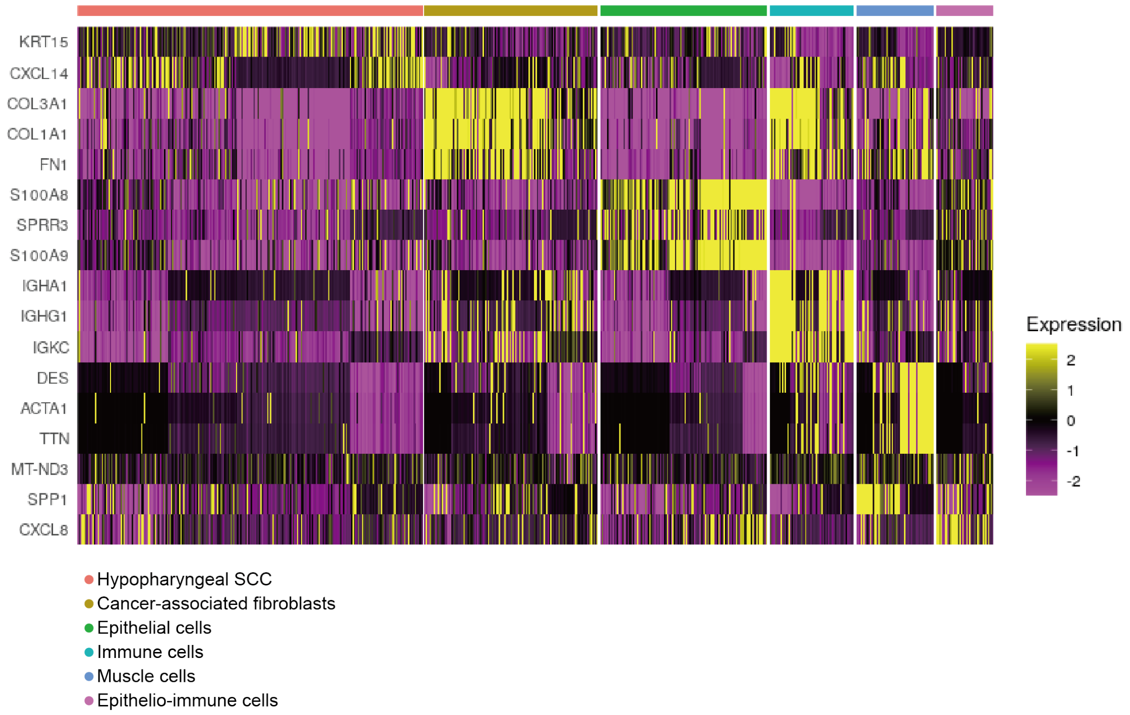


**Supplementary Figure 1.** Heatmap representation of marker gene expression in each spot cluster derived from ST data. Each row corresponds to a specific gene, while each column corresponds to a cluster within hypophayngeal SCC tissue samples. Gene expression levels are depicted by color intensity ranging from magenta (low expression) to yellow (high expression). Clusters are annotated based on the key gene expression identifiers.

**Abbreviation:** SCC, squamous cell carcinoma; ST, spatial transcriptomics

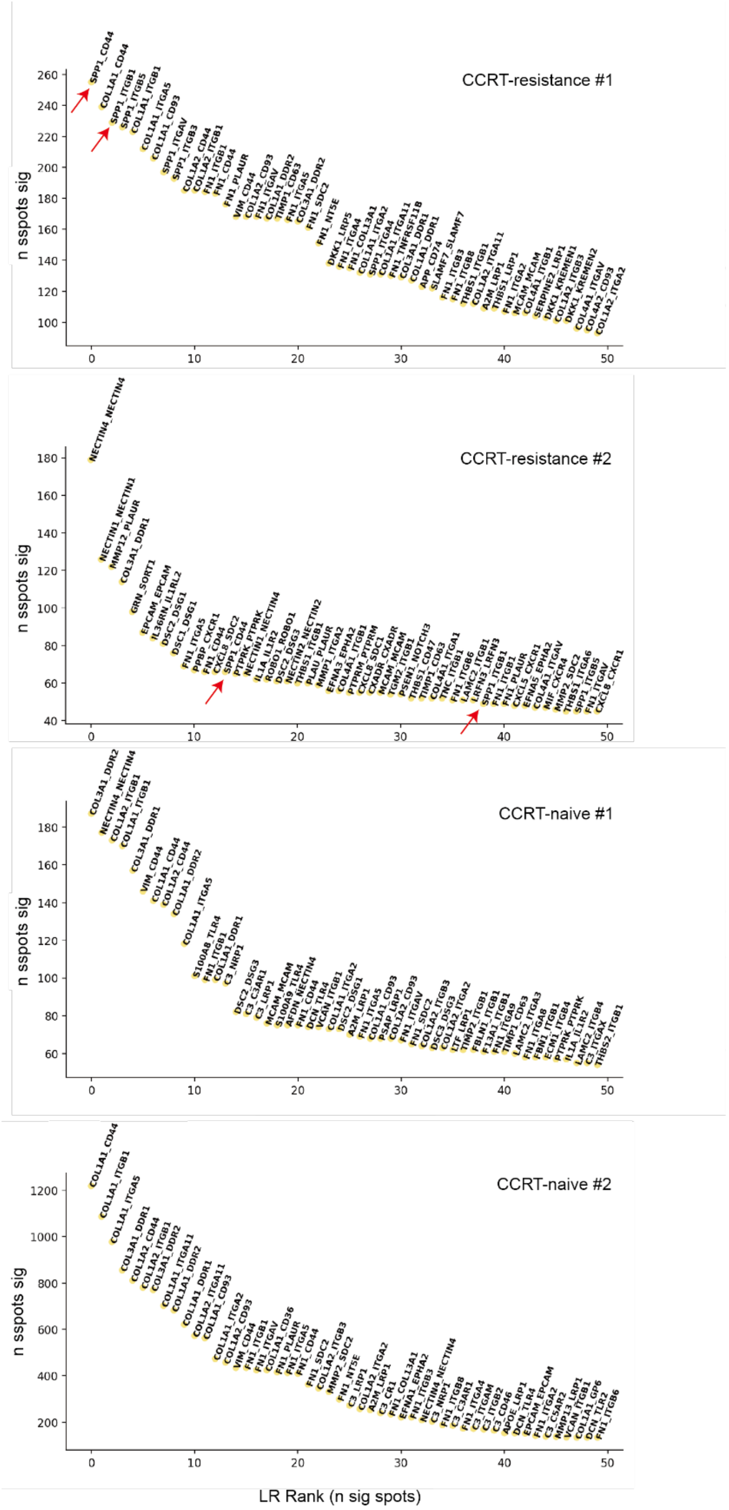


**Supplementary Figure 2.** The top 50 ligand-receptor interaction pairs were identified using the *stLearn* tool, which incorporates spatial location and co-expression in the ST data to infer the number of spots with significant interactions. In CCRT-resistance samples, *SPP1* showed a high interaction with *CD44* and *ITGB1* (indicated by red arrow).

**Abbreviation:** CCRT, concurrent chemo-radiotherapy; ST, spatial transcriptome

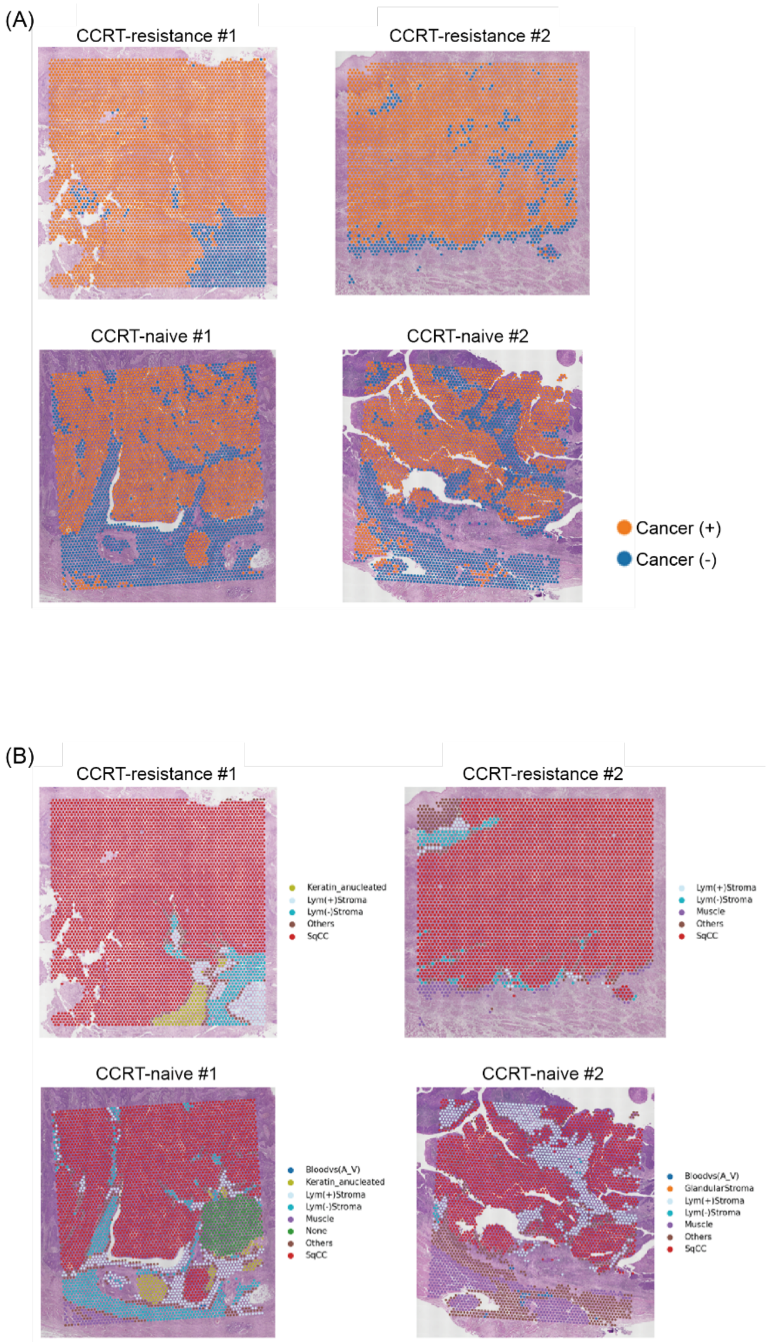


**Supplementary Figure 3.** Annotation of cancerous and non-cancerous regions. **(A)** Application of the *Cancer-Finder* algorithm to four hypopharyngeal SCC ST datasets **(B)** Annotation by a board-certified pathologist. The annotated spots were overlaid on the H&E-stained slides (refer to **Annotation of the cancer region and calculation of distance from the cancer** in **Supplementary Methods**)

**Abbreviation**: SCC, squamous cell carcinoma; ST, spatial transcriptomics

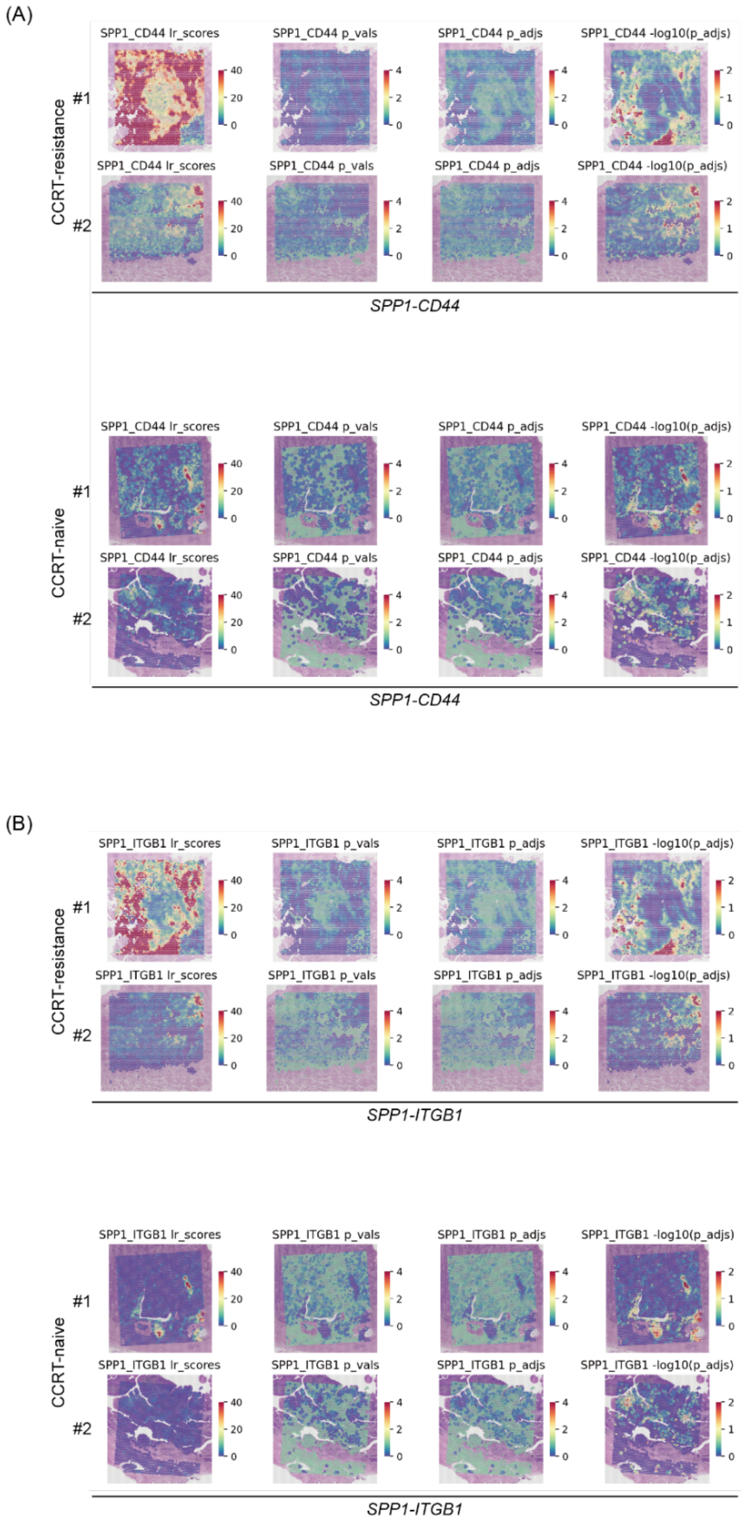


**Supplementary Figure 4.** Spatial heatmap for indexes of ligand-receptor interactions for *SPP1-CD44* **(A)** and *SPP1-ITGB1* **(B),** overlaid on an H&E-stained slide image in each sample. "lr_scores" represent the strength of ligand-receptor colocalization, "p_vals" indicate the significance of "lr_scores" determined by creating random distribution of non-interacting gene-gene pairs, and "p_adjs" show the adjusted p values.

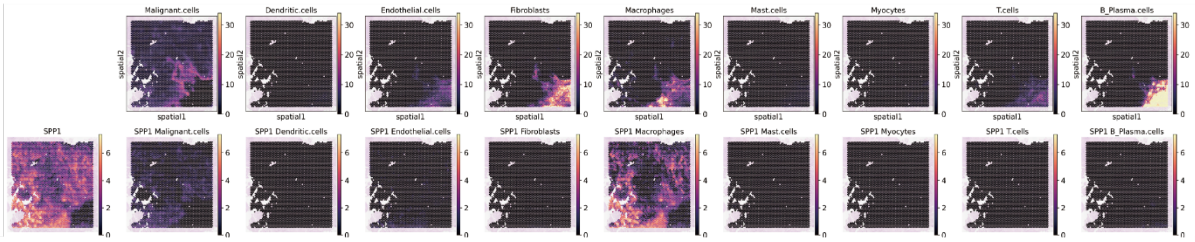


CCRT-resistance #1

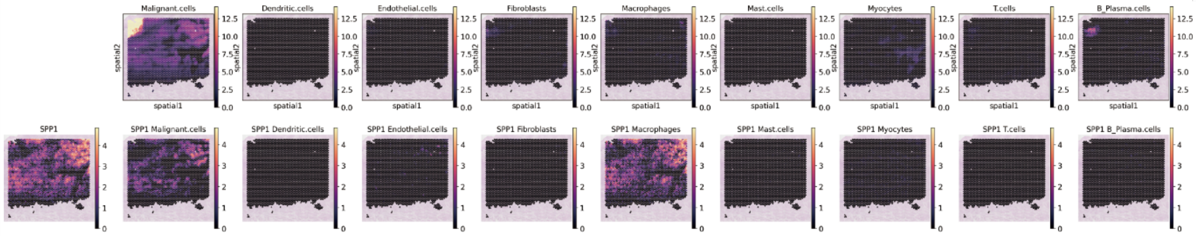


CCRT-resistance #2

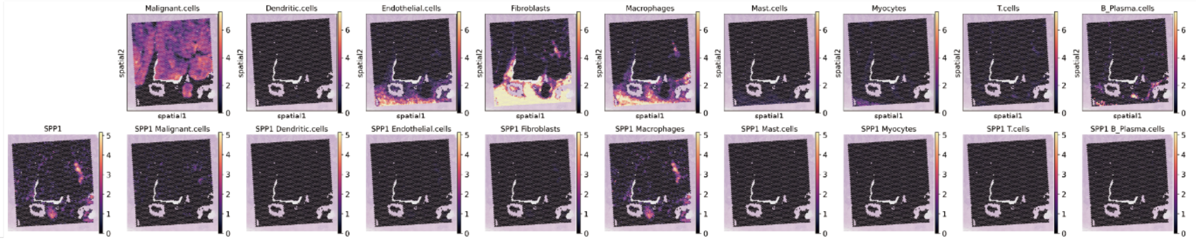


CCRT-naive #1

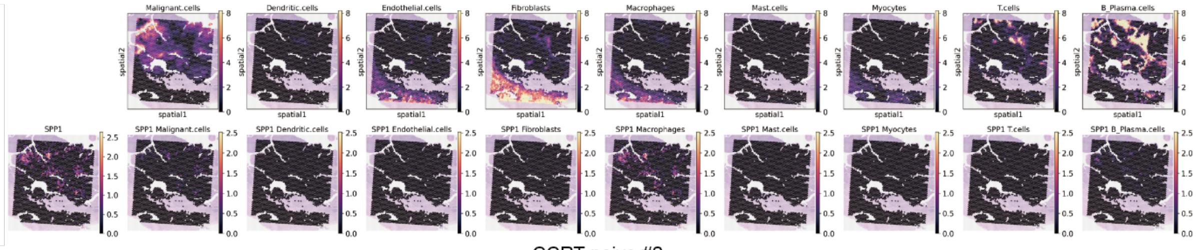


CCRT-naive #2

**Supplementary Figure 5.** Heatmaps showing the abundance of malignant epithelial cells and other cell types comprising the TME (dendritic cells, endothelial cells, fibroblasts, macrophages, mast cells, myocytes, T cells, and plasma cells) on ST spots for each hypopharyngeal SCC sample (top row of each panel). The bottom row of each panel illustrates the spatial expression of *SPP1* and cell type-specific *SPP1* expression in four hypopharyngeal SCC samples. This figure was generated using the *Cell2location* package.

**Abbreviation**: CCRT, concurrent chemo-radiotherapy**;** SCC, squamous cell carcinoma; ST, spatial transcriptomics; TME, tumor microenvironment

(A)

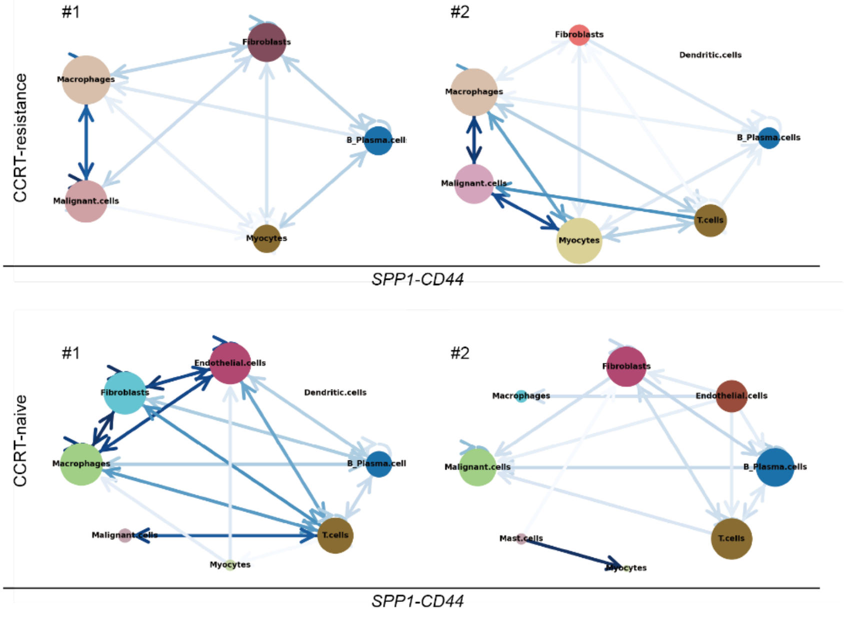


(B)

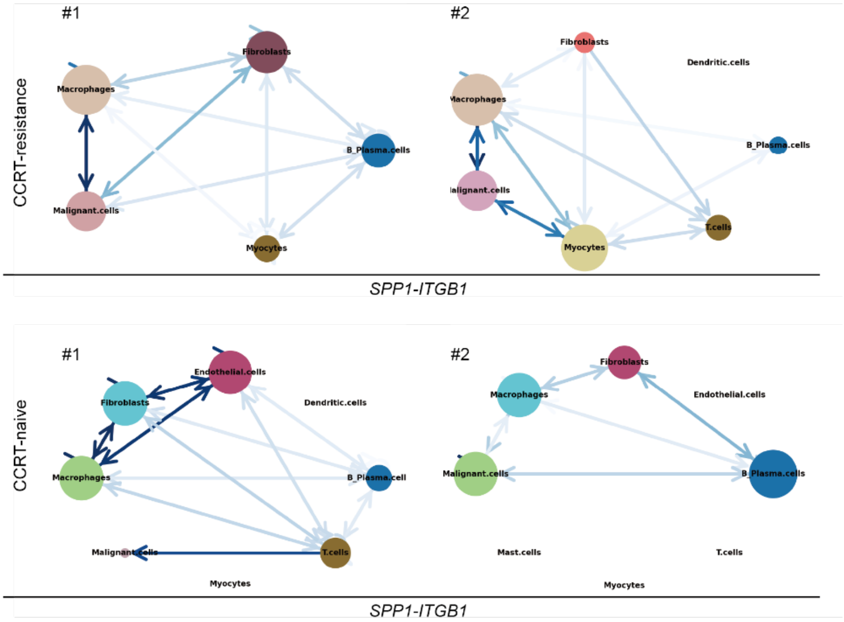


**Supplementary Figure 6.** In each study samples, cell-cell interactions were analyzed to identify predominant cells involved in ligand-receptor interactions, *SPP1-CD44* **(A)** and *SPP1-ITGB1* **(B).** The size of the spots represents the total number of significant spot interactions the cell type is involved in, and the color of the edge represents the number of significant interactions between the two cell types. In CCRT-resistance samples, *SPP1-CD44* and *SPP1-ITGB1* interactions occur primarily between malignant epithelial cells and macrophages, while these interactions occur mainly between macrophages and endothelial cells or fibroblasts in CCRT-naive samples.

**Abbreviation:** CCRT, concurrent chemo-radiotherapy

**Supplementary Methods**

**Patient enrollment**

The patients with locally advanced hypopharyngeal squamous cell carcinoma (SCC) were retrospectively enrolled from the cancer tissue bank. The clinical study was approved by institutional review board of Seoul National University Hospital (approval date: 27/12/2023, approval number: H-2203-011-1304) as a minimal-risk retrospective study and the individual consent was waived. A total of four patients were enrolled and classified into two groups. One of the groups consists of two patients who underwent upfront surgical treatment of hypopharyngeal SCC without previous concurrent chemo-radiotherapy (CCRT), named as 'CCRT-naive', The other group, named 'CCRT-resistance', also consists of two patients who had had definite CCRT as a primary treatment and underwent salvage operation for recurrence of hypopharyngeal SCC. Detailed clinical information of all study patients was shown in **Supplementary Table**.

**Sample acquisition and sequencing: Visium spatial transcriptomics (ST)**

Tumor tissues were obtained from the surgical specimen, fixed with formalin, and embedded into paraffin blocks for preservation. To process formalin-fixed, paraffin-embedded samples for RNA sequencing, the paraffin blocks were sectioned, mounted on glass slides, and deparaffinized. The tissue sections were stained with hematoxylin-eosin (H&E) and de-crosslinked. After the hybridization of the probes that capture the specific sequence of RNA, the Visium CytAssist device (10x Genomics, USA) was utilized to transfer the tissue onto the Visium slide. The tissue was permeabilized, RNAs were captured, reverse-transcribed into cDNA, and amplified to construct gene expression libraries. The libraries were sequenced, and the raw sequence files were processed from BCL to FASTQ format and then to count matrices using Space Ranger (10x Genomics, USA) custom pipeline. H&E-stained images were aligned with the RNA capture plane and processed for downstream analysis.

**Preprocessing ST datasets**

To remove the low-quality data in Visium ST, spots with a number of genes with nonzero expression greater than 500 and a percentage of mitochondrial genes lower than 20% were selected. The raw count matrix for each Visium slide was normalized and scaled using *SCTransform* [1]. *SCTransform* transforms the raw count of the gene to Pearson residuals based on regularized negative binomial regression, with the total count and percentage of mitochondrial genes as covariates. Canonical correlation analysis (CCA) was performed to embed spots from pairs of slides into low dimensional space, and mutual nearest neighbors (MNN) were found between the slides to correct the batch effect [2]. After batch correction, the dimensionality was reduced using principal component analysis (PCA), and 30 dimensions (PC1 to PC30) were chosen for the clustering of spots using Louvain algorithm (resolution = 0.15) [3]. The markers of each spot cluster were discovered using Wilcoxon rank-sum test, and the Bonferroni method was applied for multiple comparison corrections. The spot clusters were named according to the signature gene sets composed of the top 20 markers (**Supplementary File**).

All of the above analyses were performed in *Seurat* (version 4.4.0) running on R [4] and *scanpy* (version 1.10.1) running on Python [5].

**Estimation of cell type distribution in the tumor microenvironment**

The abundance of cell in each spot was estimated using *Cell2location* (version 0.1.4) [6]. *Cell2location* uses reference single-cell transcriptomics dataset and applies a negative binomial regression model to extract gene expression profiles of cell types in the tumor. A publicly available single-cell dataset obtained from 20 hypopharyngeal SCCs (accession number GSE181919) and cell type annotation information were used to infer the cell type signatures of nine cell types: cancer cells, endothelial cells, fibroblasts, myocytes, macrophages, dendritic cells, mast cells, plasma cells, and T cells [7]. Then, leveraging cell type signatures, the spatial abundance of cell types and cell type-specific expression profiles at each spot location were predicted. The number of cells per spot and detection alpha were set to 10 and 20, respectively.

**Annotation of the cancer region and calculation of distance from the cancer**

Pathological annotation of the dataset was performed by a board-certified pathologist. The tumor tissues were divided into nine categories: ‘SqCC’, Cancer regions, including invasive carcinoma and carcinoma *in situ*; ‘Lym (+) stroma’, stroma with immune cell-abundant; ‘Lym (-) stroma’, immune cell-depleted stroma; ‘Glandular stroma’, gland-rich regions; ‘Blood vs’, blood vessels; ‘Anucleated keratin’, regions with keratin materials; ‘Normal mucosa’, normal squamous mucosa; ‘Muscle’, smooth muscle regions; and ‘Necrosis’, necrotic regions. Notably, CCRT-resistance samples had low quality of the H&E stain, and the pathologic annotation was performed at the level of differentiating tumoral and stromal compartments.

To complement the pathological annotation for the low-quality H&E images and double-check the cancer region within the slide, we applied a deep learning model, *Cancer-Finder* [8]. *Cancer-Finder* was trained on ST datasets of several cancer types and classifies the spot into non-cancer and cancer spots based on transcriptomic profiles. After the segmentation of the cancer region, the annotation results were compared between those from pathologists and from transcriptomic profiles.

The physical distance from the annotated cancer region to a certain spot was defined using the function *annotation_coordinate* in the package *tacco* (version 0.01-post2) [9]. The distance between the spots was calculated by assuming the spots are placed on the hexagonal coordinates and by setting the size of the bin to 1. The estimated distance was used to evaluate the spatial patterns of genes, cell types and ligand-receptor interactions with respect to the cancer region.

**Ligand-receptor interaction analysis**

The spatial co-expression pattern between ligand and receptor (LR) was captured using a method called *stLearn* (version 0.4.12) [10]. The strength of LR interaction in the Visium slide was calculated by averaging the mean ligand expression in the spot and its surrounding neighbors expressing receptor and vice versa. The scores were calculated for all LR pairs in Omnipath database, and the top LR pairs were ranked [11]. The significance of the interaction was computed based on a random background distribution derived from 10,000 non-interacting gene pairs with similar expression levels. In addition, using the cell type abundance estimated by *Cell2location*, the cell type-specific interaction patterns were investigated. The spots were annotated by the most abundant cell type in each location, and the number of neighboring spots that express the ligand or receptor was counted. The cell labels were permutated 500 times while maintaining the cell type proportion in the whole slide to create random distribution. Then, the probability that a certain LR interaction occurs between cell type pairs was calculated. The resulting LR interactions between cell type pairs were presented with cell-cell interaction network plot.

**Supplementary File**

Top 20 gene markers for the annotation of each spot cluster.
